# Neonicotinoid exposure affects foraging, nesting, and reproductive success of ground-nesting solitary bees

**DOI:** 10.1101/2020.10.07.330605

**Authors:** D. Susan Willis Chan, Nigel E. Raine

**Affiliations:** School of Environmental Sciences, University of Guelph, Guelph, Ontario, N1G 2W1, Canada

## Abstract

Despite their indispensable role in food production^1,2^, insect pollinators are threatened by multiple environmental stressors, including pesticide exposure^2-4^. Although honeybees are important, most pollinating insect species are wild, solitary, ground-nesting bees^1,4-6^ that are inadequately represented by honeybee-centric regulatory pesticide risk assessment frameworks^7,8^. Here, for the first time, we evaluate the effects of realistic exposure to systemic insecticides (imidacloprid, thiamethoxam or chlorantraniliprole) on a ground-nesting bee species in a semi-field experiment. Hoary squash bees (*Eucera* (*Peponapis*) *pruinosa*) provide essential pollination services to North American pumpkin and squash crops^9-14^ and commonly nest within cropping areas^10^, placing them at risk of exposure to pesticides in soil^8,10^, nectar and pollen^15,16^. Hoary squash bees exposed to an imidacloprid-treated crop initiated 85% fewer nests, left 84% more pollen unharvested, and produced 89% fewer offspring than untreated controls. We found no measurable impact on squash bees from exposure to thiamethoxam- or chlorantraniliprole-treated crops. Our results demonstrate important sublethal effects of field-realistic exposure to a soil-applied neonicotinoid (imidacloprid) on the behaviour and reproductive success of a ground-nesting solitary bee. To prevent potential declines in ground-nesting bee populations and associated impoverishment of crop pollination services, soil must be considered a possible route of pesticide exposure for bees, and restrictions on soil-applied insecticides may be justified.

## Main text

Conserving pollinator biodiversity is becoming increasingly important with rising human demand for insect-pollinated crops^1,2,17-19^. Associated losses of pollination services will be particularly detrimental for growers of crops like pumpkin, squash, and gourds (*Cucurbita* crops) that are entirely dependent upon pollination by bees to set fruit^20,21^.

Among the most efficient and widely distributed wild pollinator of *Cucurbita* crops in North America is the solitary hoary squash bee (*Eucera* (*Peponapis*) *pruinosa* (Say, 1837))^11-12,21-24^. This species exhibits an unusually high degree of dietary specialization, depending entirely upon *Cucurbita* crops (Cucurbitaceae) for pollen across most of its range^22,23,25^. Each hoary squash bee female constructs its own nest in the ground^26,27^, commonly within cropping areas^10^ where systemic insecticides are often applied as soil drenches or seed treatments^8^.

Historically almost all information on impacts of pesticide exposure for bees has come from honeybees (*Apis mellifera* Linnaeus, 1758), the model species for insect pollinator pesticide risk assessments^4,7,28,29^. More recently, exposure to field-realistic levels of systemic neonicotinoid insecticides have been shown to have adverse effects on learning and memory^30,31^, foraging behaviour^32-35^, colony establishment^36,37^, reproductive success^38-40^, and the delivery of pollination services^41,42^ in bumblebees (*Bombus* spp.).

Although non-social bees have received less attention, adverse effects from exposure to neonicotinoids on nest establishment^40^, homing ability^43^, reproductive output^44^, developmental delays^45^, and reduced adult size and longevity^45^ have been demonstrated for cavity-nesting solitary bees.

While all bees that forage on treated crops may be at risk of exposure to systemic pesticide residues in pollen and nectar^28,29^, there is an additional risk of exposure to residues in soil for ground-nesting bees^4,8^. This route of pesticide exposure is currently not considered in regulatory environmental risk assessments for pollinators because honeybees rarely come into direct contact with soil^3,4,7,8^. Although approximately 70% of solitary bee species nest in the ground^4,46^ where they spend most of their lives^8,46^, and many of these are associated with agriculture^1,4-6,21,23^, no studies have yet evaluated potential impacts of pesticide exposure on this highly significant group of insect pollinators.

We are the first to evaluate the potential effects of exposure to a crop treated with one of three different systemic insecticides (neonicotinoids imidacloprid or thiamethoxam or the anthranilic diamide chlorantraniliprole) on nest establishment, foraging behaviour and offspring production for a solitary, ground-nesting bee in a semi-field hoop house study.

To assess the impacts on wild squash bees of field realistic systemic insecticide exposure we grew squash plants (*Cucurbita pepo*) in twelve netted hoop houses. We applied Admire® (imidacloprid) as a soil drench at seeding to three hoop houses; planted FarMoreFI400® (thiamethoxam + fungicides: Cruiser 5FS® + Apron XL®, Maxim 4FS® and Dynasty®) treated seeds in three more hoop houses; applied Coragen® (chlorantraniliprole) as a foliar spray to another three hoop houses, with the remaining three hoop houses left untreated as insecticide controls (Methods; Extended Data; Fig. 1).

**Figure 1.**
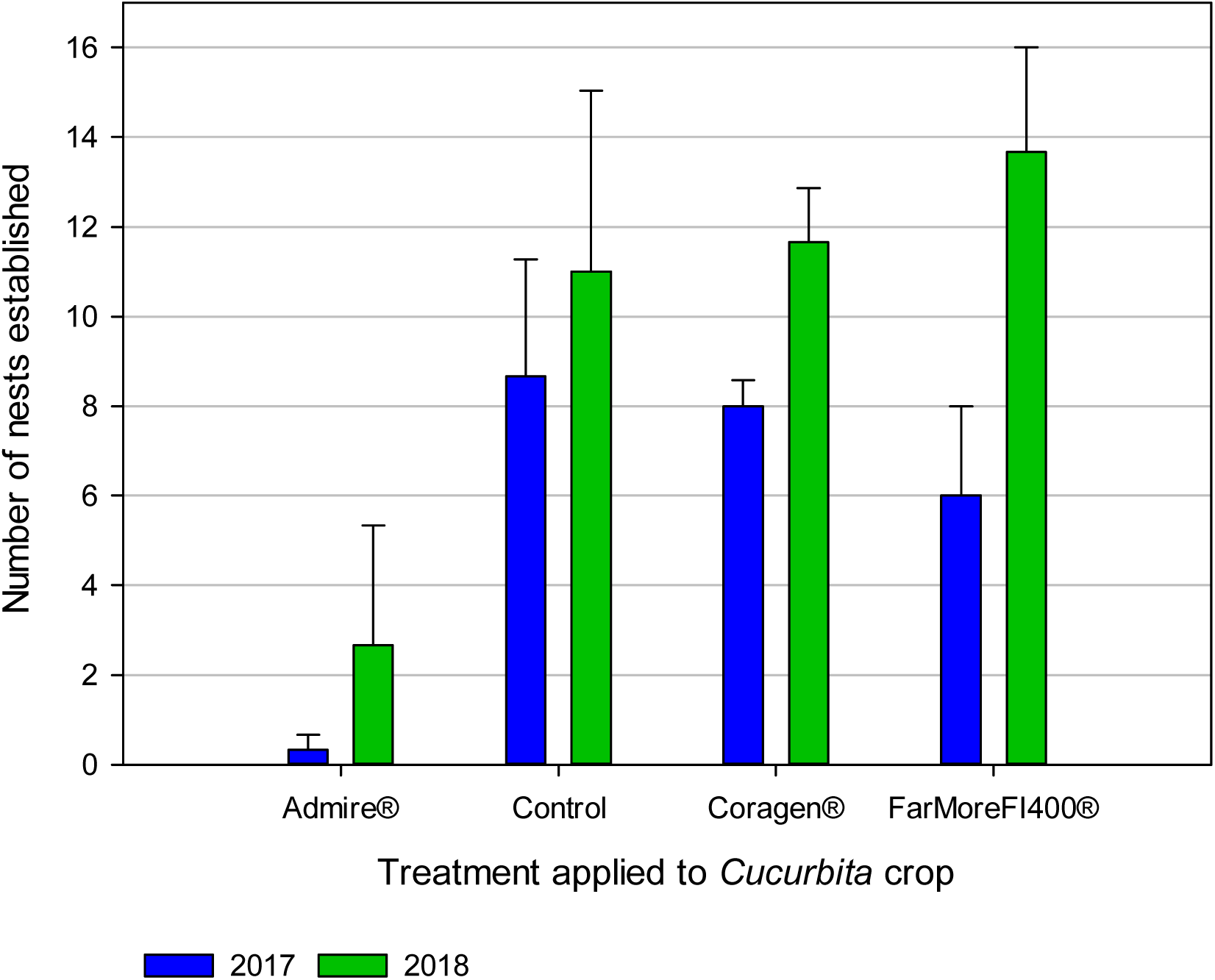
Numbers of hoary squash bee (*Eucera* (*Peponapis*) *pruinosa*) nests constructed over the 2017 and 2018 seasons in hoop houses in which one systemic insecticide treatment (either Admire®-imidacloprid, applied to soil at seeding; or Coragen®-chlorantraniliprole applied as a foliar spray; or FarMoreFI400®-thiamethoxam applied as a seed treatment) was applied to the *Cucurbita* acorn squash (Varieties: TableStar® (2017), Celebration® (2018)) crop 8-weeks before the bee active period, or the crop remained untreated as a control. Data presented are means ± SE across three hoop houses per treatment in each year.

We introduced mated female hoary squash bees into each hoop house at crop bloom (i.e., 8-weeks later: Methods). After mating, hoary squash bee females begin their nesting phase involving three critical overlapping activities: (1) soil excavation and nest establishment; (2) foraging for pollen and nectar to provision nest cells; and (3) laying eggs^26,27^.

To determine if exposure to squash crops treated with systemic insecticides interfered with nest establishment, we tracked the number of active nests made by foundress (2017) and second generation (2018) females in each hoop house (Methods). Nest establishment was substantially affected by insecticide treatment (F_3,14_ = 7.33; p = 0.0034; Table S1; Table S2), with significantly fewer nests established in the Admire®-treated hoop houses than in other treatments (mean ± SE: Admire®: 1.50 ± 1.31; Control: 9.83 ± 2.21; Coragen®: 9.83 ± 1.01; FarMoreF1400®: 9.83 ± 2.20; Fig. 1; Table 1). Compared to the untreated control, nest establishment by hoary squash bees exposed to Admire®-treated squash crops was reduced by 76% in 2017 (mean ± SE: Admire®: 0.33 ± 0.33; Control: 8.67 ± 2.60), 96% in 2018 (Admire®: 2.67 ± 2.67; Control: 11.00 ± 4.04) with an 85% reduction across both years (Admire® vs Control: t_13_ = -3.83, p = 0.0088; Fig. 1; Table 1; Table S3). We detected no significant differences in the numbers of nests established by females in the control, Coragen®, and FarMoreF1400® treatments (Table 1; Table S3).

**Table 1.** Summary statistics (means ± standard error) for all dependent variables and insecticide treatments in each year and combined for both years.

| Dependent Variable | Treatment | 2017 | 2018 | Combined |
| --- | --- | --- | --- | --- |
| Total Nests | Admire® | 0.33 ± 0.33 | 2.67 ± 2.67 | 1.50 ± 1.31<sup>a</sup> |
| | Control | 8.67 ± 2.60 | 11.00 ± 4.04 | 9.83 ± 2.21<sup>b</sup> |
| | Coragen® | 8.00 ± 0.58 | 11.67 ± 1.20 | 9.83 ± 1.01<sup>b</sup> |
| | FarMore F1400® | 6.00 ± 2.00 | 13.67 ± 2.33 | 9.83 ± 2.20<sup>b</sup> |
| Unharvested Pollen | Admire® | 13662.0 ± 1179.6<sup>a</sup> | | |
| | Control | 2168.0 ± 461.5<sup>b</sup> | | |
| | Coragen® | 5422.0 ± 1075.4<sup>c</sup> | | |
| | FarMore F1400® | 2606.0 ± 457.0<sup>bc</sup> | | |
| Total Offspring | Admire® | 3.33 ± 1.45<sup>a</sup> | 3.33 ± 3.33<sup>a</sup> | 3.33 ± 1.63<sup>a</sup> |
| | Control | 23.00 ± 10.69<sup>b</sup> | 42.00 ± 8.19<sup>b</sup> | 35.00 ± 8.30<sup>b</sup> |
| | Coragen® | 18.00 ± 3.21<sup>ab</sup> | 42.67 ± 13.37<sup>b</sup> | 32.33 ± 9.07<sup>b</sup> |
| | FarMore F1400® | 21.67 ± 4.18<sup>ab</sup> | 20.00 ± 7.09<sup>b*</sup> | 22.17 ± 4.69<sup>b</sup> |
| Total Flowers | Admire® | 45.44 ± 3.40 | 54.42 ± 0.87 | 49.93 ± 2.55 |
| | Control | 42.83 ± 8.22 | 46.46 ± 3.88 | 44.65 ± 4.15 |
| | Coragen® | 48.06 ± 4.84 | 45.42 ± 7.30 | 46.74 ± 3.96 |
| | FarMore F1400® | 54.61 ± 4.57 | 40.21 ± 2.99 | 47.41 ± 4.04 |
| Sex Ratio | Admire® | 0.5<sup>†</sup> | 4<sup>†</sup> | 1.67 ± 1.67<sup>a</sup> |
| | Control | 1.17 ± 0.31 | 2.27 ± 0.46 | 1.59 ± 0.27<sup>a</sup> |
| | Coragen® | 0.51 ± 0.18 | 4.42 ± 3.42 | 2.01 ± 1.31<sup>a</sup> |
| | FarMore F1400® | 0.89 ± 0.29 | 3.89 ± 1.84 | 1.73 ± 0.67<sup>a</sup> |
| % Fruit Set | Admire® | 55.92 ± 11.30<sup>a</sup> | | |
| | Control | 54.60 ± 7.80<sup>a</sup> | | |
| | Coragen® | 45.83 ± 4.16<sup>a</sup> | | |
| | FarMore F1400® | 54.17 ± 17.81<sup>a</sup> | | |
| % Marketable Fruit (>500 g) | Admire® | 67.46 ± 9.58<sup>a</sup> | 76.46 ± 3.33<sup>a</sup> | |
| | Control | 61.93 ± 9.30<sup>a</sup> | 84.54 ± 2.10<sup>a</sup> | |
| | Coragen® | 62.56 ± 4.11<sup>a</sup> | 76.73 ± 9.08<sup>a</sup> | |
| | FarMore F1400® | 68.37 ± 4.24<sup>a</sup> | 70.77 ± 3.68<sup>a</sup> | |

\* Significant difference is marginal (p = 0.0585).
<sup>†</sup> Standard error could not be calculated for the sex ratio of Admire® in 2017 and 2018 because bees only emerged in one hoop house in each year.

These results are consistent with studies indicating neonicotinoid-exposure can reduce nesting activity in stem-nesting solitary bees^40,44^ and delay or reduce the likelihood of nest establishment in bumblebees^36,37,47^. Our study is the first to report an impact of neonicotinoid-exposure on nesting in any solitary ground-nesting species, arguably the most ecologically important and biodiverse group of bees. The 85% reduction in nest establishment (over 2 years) by squash bee females exposed to imidacloprid under real agricultural conditions incorporates both contact with soil and consumption of contaminated nectar and pollen as potential routes of exposure, whilst previous studies have focused solely on oral exposure^36,37,40,44,47^.

During their nesting phase, hoary squash bees also amass pollen to provision their nest cells. The reproductive success of female solitary bees is limited by the number of nest cells they can provision^44,46^, and the amount of pollen harvested affects the subsequent reproductive success of their offspring^48-50^.

To determine if exposure to crops treated with systemic insecticides affected pollen harvesting by female hoary squash bees, we counted the pollen remaining on anthers of squash flowers at the end of the daily foraging period in each hoop house (Methods; Extended Data Table 1). Pollen collection was substantially affected by insecticide treatment (F_3,114_ = 37.82; p < 0.0001; Table S1; Table S2) with 84% more unharvested pollen remaining on anthers in the Admire®-treatment than the control (Admire®: 13662.0 ± 1179.6; Control: 2168.0 ± 461.5; t_114_ = 9.38, p < 0.0001; Fig. 2; Table S3). Although hoary squash bees in the Coragen® treatment also collected significantly less pollen than bees in the control group (Coragen®: 5422.0 ± 1075.4; Control: 2168.0 ± 461.5; t_114_ = -2.65, p = 0.0443; Table 1; Table S3), nest establishment appeared unaffected (Table S2). There was no significant difference in the quantity of pollen collected by bees in the FarMoreF1400® (2606.0 ± 457.0) and the control treatment (t_114_ = -0.36, p = 0.9843; Table 1; Table S3).

**Figure 2.**
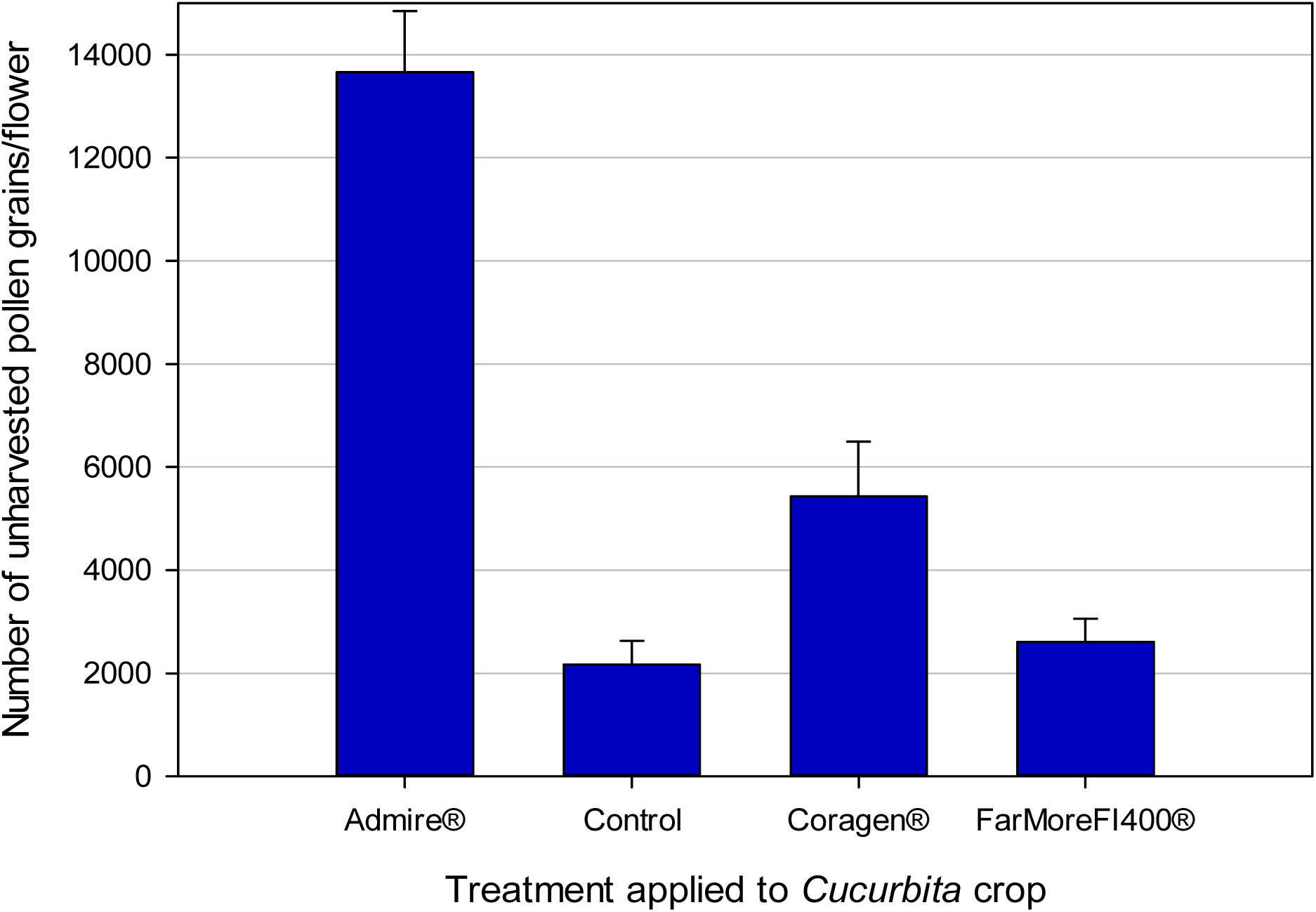
Number of pollen grains remaining unharvested on each squash flower (*Cucurbita pepo*, var. TableStar®) anther at the end of the squash bee (*Eucera* (*Peponapis*) *pruinosa*) daily foraging period (dawn to 11 am). Bees were kept in hoop houses in which one systemic insecticide treatment (either Admire®-imidacloprid, applied to soil at seeding; or Coragen®-chlorantraniliprole applied as a foliar spray; or FarMoreFI400®-thiamethoxam applied as a seed treatment) was applied to the crop 8-weeks before the bee active period, or the crop remained untreated as a control. Data presented are means ± SE across three hoop houses per treatment from 2017.

Our results show that female hoary squash bees exposed to Admire®-treated crops constructed fewer nests and collected less pollen than unexposed bees, a combined negative effect that would likely reduce offspring production^48-50^. To determine if offspring production was affected by exposure to crops treated with systemic insecticides, we counted and sexed all hoary squash bees that emerged from all nests constructed in the hoop houses in 2017 and 2018 (Methods). Offspring production was significantly affected by insecticide treatment (F_3,13_ = 14.19, p = 0.0002), year (F_1,13_ = 11.79, p = 0.0044) and the abundance of flowers (F_1,13_ = 10.44, p = 0.0066) as main model effects, and the interaction between insecticide treatment and year (F_3,13_ = 3.81, p =0.0368; Table S1; Table S2). In the Admire®-treatment, 89% fewer offspring were produced than in the control (Admire® = 3.33 ± 1.63, Control = 35.00 ± 8.30; t_15_ = -5.98, p = 0.0002; Table 1; Table S3).

Although ∼36% fewer offspring were produced in the FarMoreF1400® treatment than in the control (FarMoreF1400® = 22.17 ± 4.69; Table 1), no significant difference was detected (t_15_ = 2.59, p = 0.0910; Table S3). There was also no significant difference in offspring produced between the Coragen® treatment (32.33 ± 9.07; Table 1) and the untreated control (t_15_ = 0.80, p = 0.8542; Table S3). The effect of exposure to FarMoreF1400®, though not statistically significant, may warrant more attention as clothianidin, a metabolite of thiamethoxam^51,52^ can have toxic effects on honeybees and can also affect stem-nesting solitary bees^40,44,53^.

Floral resources in the hoophouses substantially affected the number of offspring produced (F_3,13_ = 10.44; p = 0.0066; Table S2) but did not explain the variance in the number of offspring well (Y = 1.157 + 0.467X; r^2^ = 0.0423). The effects of year, and the treatment*year interaction, are likely explained by expanding populations in the second generation (2018) in the Coragen® and control hoop houses but not the Admire® and FarMore F1400® treatments (Table S2; Table S4).

While cavity-nesting solitary bees can shift towards male-biased offspring sex ratios after exposure to sublethal doses of neonicotinoids^44^, we found no effect of exposure to systemic insecticide treatments (Admire®, Coragen®, FarMoreF1400®) on the sex ratio of hoary squash bees here (Model: Sex Ratio = Treatment + Year ; Treatment: F_3,14_ = 0.08, p = 0.9679; Year: F_1,14_ = 6.73, p = 0.0212; Table S1; Table S2). Offspring sex ratios varied substantially between years within all treatments, becoming more male biased overall between 2017 and 2018 (2017: 0.79 ± 0.14; 2018: 3.52 ± 1.05; Table 1). It is unlikely that the shift towards more male biased sex ratios in 2018 was the result is of weather conditions such as flooding or severe cold, which should have impacted shallower (male-bearing) nest cells more than deeper (female-bearing) nest cells^46,54^. Alternatively, as hoop house populations expanded from 8 to ∼16.5 bees from 2017 and 2018, increased competition among females for floral resources may have caused them to allocate limited pollen resources to the production of male offspring that require less pollen to develop^55^.

For all nesting phase effects (i.e. nest establishment, pollen collection, and offspring production), there is strong evidence of negative impacts of exposure to an Admire®-treated crop under ecologically and agriculturally realistic field conditions in our study. There may also be negative impacts on offspring production from exposure to FarMoreFI400® which we were unable to detect statistically.

Potential routes of exposure to systemic insecticides here include topical exposure to soil-applied neonicotinoids (Admire® applied in-furrow at seeding or FarMoreFI400® applied as seed coating) during nest construction, topical exposure to chlorantraniliprole applied to crop leaves, and topical and oral exposure to all three systemic insecticides in the nectar and pollen of the squash crop. Which route(s) contributed most to the observed effects could not be discriminated, representing an important knowledge gap for future research.

For hoary squash bees exposed to an Admire®-treated squash crop, the number of nests established was strongly reduced (Fig. 1), less pollen was collected (Fig. 2) and fewer offspring were produced (Fig. 3). These outcomes for Admire®-exposed hoary squash bees are substantial and should be of concern to growers, pesticide regulators, and those engaged in pollinator conservation.

**Figure 3.**
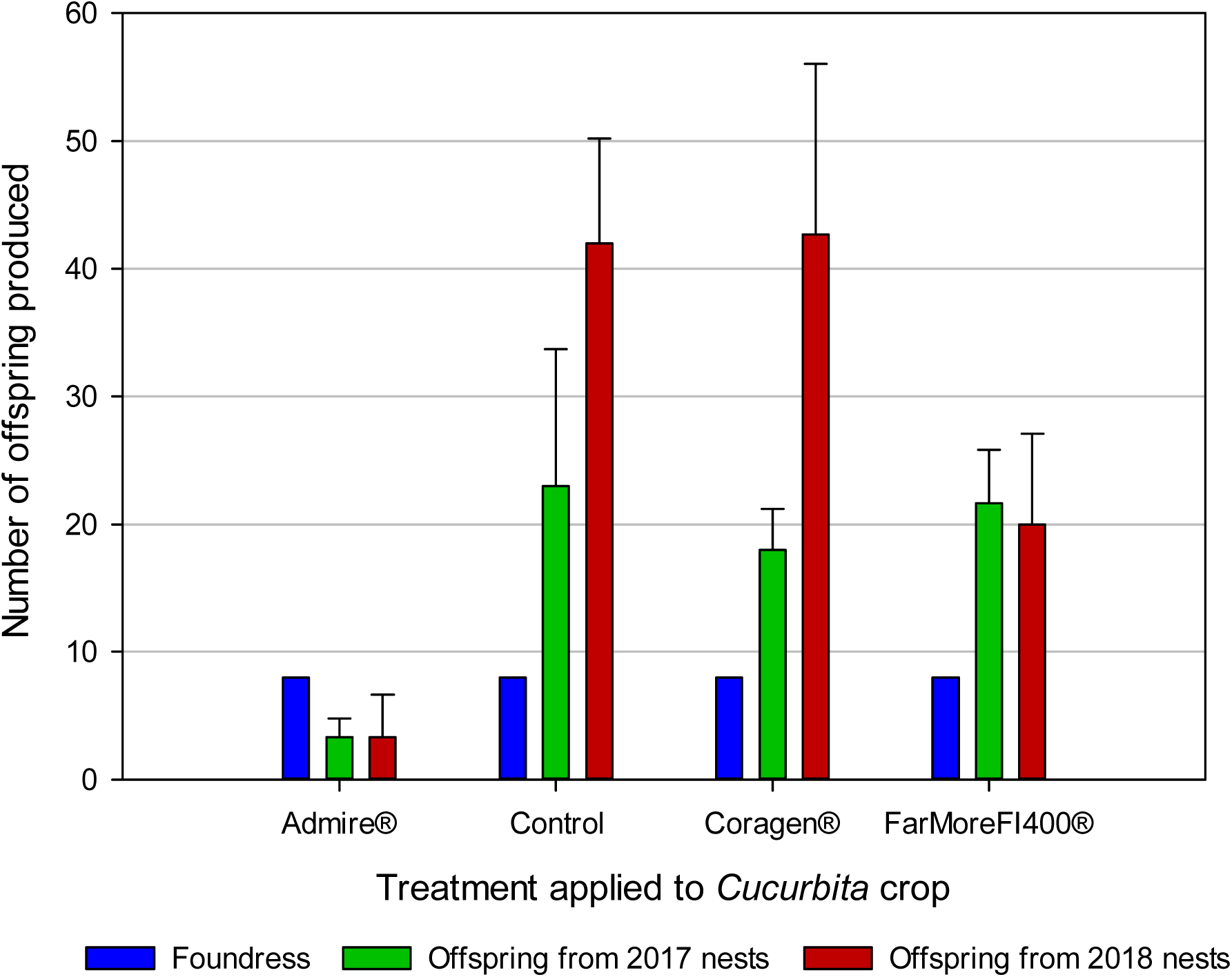
Number of hoary squash bee offspring from nests established by mated female bees in 2017 and 2018. In 2017, eight mated foundress hoary squash bee females were introduced into each hoop house in which one systemic insecticide treatment (either Admire®-imidacloprid, applied to soil at seeding; or Coragen®-chlorantraniliprole applied as a foliar spray; or FarMoreFI400®-thiamethoxam applied as a seed treatment) was applied to the squash crop (Varieties: TableStar® (2017), Celebration® (2018)) 8-weeks before bee active period, or the crop remained untreated as a control. Data presented are means ± SE across three hoop houses per treatment in each year.

For squash crops, the pollination services of bees are especially important to growers because the pollen cannot be moved between flowers by wind^20^, and the size of the pollen load deposited on the stigmas of female flowers subsequently affects fruit quality^56^. We evaluated the effect of exposure to crops treated with systemic insecticides (Admire®, Coragen®, or FarMoreF1400®) on the hoary squash bee’s ability to deliver pollination services to the crop as measured by percentage of both fruit set and marketable fruit (>500g) (Methods; Extended Data Table 1). We found no measurable effect of treatment on either fruit set (F_3,8_ = 0.16, p = 0.9186) or marketable fruit yield (F_3,18_ = 0.14, p = 0.9360; Table 1; Table S1; Table S2).

Although hoary squash bees in Admire®-treated hoop houses removed moved much less pollen from squash flowers, sufficient pollen was moved between flowers in all treatments to achieve the physiological fruit set limit of the squash crop (about 50%; Table 1)^56^. Thus, although hoary squash bees may not reproduce normally when exposed to an Admire®-treated squash crop, they may still provide effective crop pollination services within a season.

However, those services could be severely reduced in subsequent seasons unless hoary squash bees disperse into the crop from other locations, a poor long-term strategy for achieving stable pollination services for squash crops. Clearly, the issue of pollinator population health and stability would not be detected by growers if they use yield within a single season as their measure. Although replacing wild pollinators with managed honeybees might be suggested in the short term, it is not a sustainable solution because global demand for managed hives to pollinate crops has already outstripped supply^18,19,57^. Furthermore, honeybees are often not the most effective crop pollinators on an individual basis for many insect dependent crops (including for pumpkin and squash^11,12,14^), and yield and quality of many crops are improved by exposure to the pollination services of many pollinator taxa^1,2,17^.

It is notable that exposure to Coragen®-treated crops (chlorantraniliprole) seemed to have little impact on hoary squash bee reproduction in our study. As such, Coragen® may offer a viable crop protection alternative for squash growers against insect pests that poses a lower risk to ground-nesting solitary bees than some other registered systemic insecticides. The negative implications for hoary squash bees of exposure to Admire®(imidacloprid) treated crops may be an indication of a wider negative impacts on other ground-nesting solitary bee species that forage on or nest around a wide variety of other neonicotinoid-treated crops^1,5,6^. As such, the importance of our findings should resonate with regulatory agencies seeking to ensure the environmental safety of pesticides, growers who depend on the pollination services of ground-nesting bees, and conservationists seeking to understand the mechanisms for pollinator decline.

## Methods

### Study site

To establish a captive hoary squash bee population and study the potential effects of exposure to systemic insecticide treatments under controlled conditions, twelve (12) hoop houses (Width 4.80m x Length 6.10m x Height 3.05m) were set up on a farm in Peterborough County, Ontario in 2017 (Extended Data Fig. 1). All hoop houses were covered with 50% shade cloth to prevent introduced hoary squash bees from escaping and to exclude other bees from entering, while allowing exterior environmental conditions to prevail within the enclosures (Extended Data Fig. 2).

All doorways had both an outwards-opening door and a double plastic sheet across them inside to prevent bees from escaping when researchers entered or left the hoop house. Before hoary squash bees were introduced, all hoop houses were tested to ensure they were bee proof by introducing a hive of honeybees (*Apis mellifera*) and watching for escapees. Honeybees are similar in size to hoary squash bees and were unable to escape from the hoop houses, although they actively crawled over the netting. Colonies were removed from the site at night when all honeybees were back in the hive. Subsequent observations of hoary squash bees confirmed that they were not able or even trying to escape from hoop houses.

### Growing squash plants and insecticide treatments

Inside each hoop house we established two growing areas for 28 plants (14 per growing area) at spacings recommended by the seed supplier (Rupp Seeds, Inc.) and a bare soil nesting area surrounded by a mulched path (Extended Data Fig. 2). Soil at the study site was Otonabee loam, providing an excellent substrate for growing cucurbits and for hoary squash bees to construct their nests below ground. The site had a soil texture gradient of increasing sand from north to south (Extended Data Fig. 1).

Acorn squash seeds were planted into the growing areas in late May each year (varieties were Table Star® in 2017 and Celebration® in 2018), following normal Ontario farming practice. Seeds that did not germinate within five days were subsequently replanted. Untreated seeds were planted in nine hoop houses, and FarMore FI400® treated seeds (neonicotinoid insecticide: thiamethoxam) were planted in the other three (2C, 3B, 4A; Extended Data Fig. 1). Admire® (neonicotinoid insecticide: imidacloprid) and Coragen® (anthranilic diamide insecticide: chlorantraniliprole) treatments used the highest labelled rate of pesticide application following the Ontario Ministry of Agriculture, Food and Rural Affairs (OMAFRA) Vegetable Crop Protection Guide^58^, and were applied by a licensed pesticide applicator with a backpack sprayer (Roundup 190367-4 gallon), calibrated before each application.

Admire® (imidacloprid) was applied in-furrow at time of seeding (18 mL/100 m row)^58^ to three of the hoop houses (1A, 3C, 4B; Extended Data Fig. 1). Coragen® (chlorantraniliprole) was applied as a foliar spray at the 5-leaf stage of plant growth (250 mL/ha)^58^ in another three hoop houses (1C, 2B, 3A; Extended Data Fig. 1). In three more hoop houses (2C, 3B, 4A; Extended Data Fig. 1), seeds coated with FarMore FI400® Technology (Insecticide: Cruiser® 5FS insecticide = thiamethoxam + Fungicides: Apron XL® = mefanoxam, Maxim® 480FS = fludioxonil, and Dynasty® = azoxystrobin) were planted. The remaining three hoop houses (1B, 2A, 4C; Extended Data Fig. 1) were not treated with any pesticides as a control. Plants in all hoop houses were covered with row cover until the onset of flowering to provide a physical barrier against cucumber beetle (*Acalymma vittatum*) pests.

Insecticide treatments were assigned to hoop houses in a complete block design with three blocks of four treatments, and all treatments were equally represented along the site soil gradient (Extended Data Fig. 1). All treatments were assigned to the same hoop houses in both 2017 and 2018. Observations were made blind with respect to treatment in 2017, but one observer (D.S.W.C.) was aware of the treatment assignments for logistical reasons in 2018. Squash plants were also planted in the hoop houses in 2019 to provide resting sites for emerging bees, making it easier to capture them. No pesticides were applied in 2019.

Squash plants began flowering during the third week of July in all years. At the start of bloom in 2017 we discovered that the nectary of staminate (male) flowers in the Table Star® squash variety would be inaccessible to bees because these flowers lacked holes in the base of the stamen tissue covering the nectary. To overcome this issue, team members created three artificial holes in each staminate flower at the start of every day by inserting a dissection needle through the stamen tissue above the nectary (Extended Data Fig. 3). This effectively mimicked the holes that should have been present and allowed bees full access to the nectar supply throughout the experiment. We avoided this issue in 2018 by planting a different acorn squash variety (Celebration®).

### Hoary squash bee study population

To establish a population of hoary squash bees within the hoop houses, mated female hoary squash bees were captured from a wild population on a farm near Guelph, Ontario that has a large, well-established nesting aggregation with more than 3000 nests. To ensure that females to be introduced to the hoop houses were mated, only females entering a nest with a full pollen load were collected from the source population (Extended Data Fig. 4). In August 2017, we collected 96 female hoary squash bees so that we could introduce eight per hoop house. Each bee was placed in a separate aerated 2-mL microcentrifuge tube (Extended Data Fig. 5) kept upright in cool, dark conditions during transport. Upon arrival in Peterborough county, the bees were transferred to a refrigerator until the following morning when eight bees were haphazardly selected and released individually into each hoop house.

Releasing bees in the early morning coincided with the beginning of the daily bloom period of squash flowers ensuring that suitable food was immediately available. All bees (n = 96) introduced in 2017 survived the stress of capture, transport, and release into hoop houses and immediately flew out of the tubes on their own. Females began establishing nests as early as four days after introduction. To our knowledge this is the first time that mated adult hoary squash bees have been successfully introduced into and maintained under controlled semi-field conditions, offering the possibility of using hoary squash bees as a model species for other ground-nesting bees in insecticide risk assessments and research.

A second generation of hoary squash bees emerged in the hoop houses in summer 2018 from nests established by females introduced the previous summer (2017). This second generation of bees foraged, mated, and established nests in 2018. All 2018 observations were made on this second-generation and no new bees were introduced. We recorded the number and sex of all bees that emerged in the twelve hoop houses in 2018 and 2019 from nests established by the first- and second-generation females in the respective previous year.

### Nest initiation

Observations of nesting activity were made when female hoary squash bees were active, starting from 06:00 (dawn) until bee activity ceased around 11:00 each day, by four observers on ten and eight of the observation days in 2017 and 2018 respectively (Extended Data Table 1). Each observer was responsible for surveying three hoop houses per day. At the beginning of every observation day, observers searched for active nests within the hoop house and uniquely identified them with a marker (Extended Data Fig. 6). Active nests were easy to locate as bees with bright yellow pollen entering the nest were conspicuous (Extended Data Fig.4), and nests tended to be aggregated.

### Pollen collection

Female squash bees are able collect all the nutrients they require to both survive and provision their offspring from the nectar and pollen from *Cucurbita* crops^23^. Staminate (male) flowers provide both these resources, while pistillate (female) flowers produce only nectar^20^. To determine any potential variation in resource availability among treatments the total number of staminate and pistillate flowers in each hoop house was counted on six and eight of the observation days in 2017 and 2018 respectively (Extended Data Table 1). Although it was not possible to directly examine pollen loads collected by individuals without unduly disturbing the bees, pollen harvesting behaviour by the female population within a hoop house was evaluated by quantifying the number of unharvested pollen grains remaining on anthers of staminate flowers at the end of the foraging period (data were collected on 17 and 23 August 2017: Extended Data Table 1).

To do this, ten anthers were removed from staminate flowers in each hoop house and placed individually into 2-mL microcentrifuge tubes containing 0.5 mL of 70% ethanol. Pollen was dislodged from anther samples by centrifugation at 2500 rpm for three minutes and then the anther was removed. Subsequently, each microcentrifuge tube was topped up with 1.5 mL of 50% glycerine solution to bring the liquid volume to 2 mL and increase the solution viscosity^9^. Before removing aliquots, pollen samples were thoroughly agitated in the glycerine solution using a mini-vortex mixer. We calculated the number of pollen grains remaining on each anther by averaging counts of pollen grains across five 5-µL aliquots taken from each tube and relating this back to the full 2-mL volume of the pollen-glycerine-alcohol suspension.

### Offspring production

Hoary squash bees are univoltine, producing a single generation of offspring each year^27^. During the 30 to 45-day period that adults are active, females provision nest cells and lay eggs that hatch and develop to the pre-pupal stage within several weeks^27^. The bees remain at the pre-pupal stage throughout the fall, winter, and spring, pupating just before emerging as adults the following summer (July-August)^27^. Because offspring emerge as adults the year after eggs were laid into nest cells, offspring counts could only include those individuals that survived the winter and developed to maturity.

To determine the number of bees that emerged from nests established in 2017, bees in each hoop house were collected from flowers at the end of the 2018 season. This approach was taken because the hoary squash bees emerged over an extended period. All the wilted flowers in each hoop house were examined during the afternoons of August 23 and 24 and all unmated females and males found resting inside them were sexed, counted, and removed. Mated female bees entering nests or gathering pollen on flowers were collected in the same way at dawn on the mornings of August 24 and 25. As no bees were found in any hoop houses on the morning of August 25 or thereafter we concluded that offspring emergence had finished. Although this method may underestimate offspring emergence if some individuals died before the final collection, it provides useful information about the relative numbers of male and female bees that could not otherwise be quantified. Bees emerging from nests established in 2018 were collected from wilted flowers from July 30-September 4, 2019, after which no more bees emerged. After collection, each bee was sexed and released.

### Squash fruit set and marketable yield

To evaluate fruit set individual pistillate flowers were marked within each hoop house with a flag and by scratching the date and hoop house number onto the skin of the undeveloped fruit (ovary). This process was repeated on ten days (August 4-24, 2017; Extended Data Table 1) over the 3-week period during which hoary squash bees were active in the hoop houses, although pistillate flowers were not always in flower in each hoop house on any single day. At the end of the season, each ovary was evaluated as either aborted or having set fruit. Fruit set could not be evaluated in 2018 because the Celebration® variety did not tolerate the marking technique and aborted all marked fruit. Marketable yield was evaluated by harvesting, counting, and weighing all fruit from each hoop house and calculating the percentage of marketable size (≥500 g).

### Insecticide residues

To evaluate soil insecticide residues for each treatment during the period of female hoary squash bee activity, we took soil samples from the planted areas in each hoop house in 2017 and 2018 on three days during the bee active period (July and August; Extended Data Fig. 1). Samples were taken from the top 15 cm of soil because this is the depth at which hoary squash bees construct their nests^27^. For each planted area, four soil core samples were combined and subsampled to produce a single 3-g sample (i.e. 2 samples per hoop house). Potential for cross-contamination of samples was minimised by using disposable gloves, single-use containers and instruments, and thorough cleaning of the soil corer between hoop houses and a double rinse with clean water. Nectar and pollen samples were not taken within hoop houses because of concerns about depleting food resources for the study bees.

All samples were kept frozen at -20°C before submission for analysis to University of Guelph Agri-Food Laboratories (ISO/IEC 17025 accredited) which used liquid chromatography/electrospray ionization-tandem mass spectrometry (LC/ESI-MS/MS) and gas chromatography-tandem mass spectrometry (GC-MS/MS) (Modified Canadian Food Inspection Agency (CFIA) PMR-006-V1.0) to detect the presence and determine the concentrations of imidacloprid, clothianidin, thiamethoxam, and chlorantraniliprole in samples. Although clothianidin was not applied as a treatment, it is a common, persistent metabolite of thiamethoxam^51^ and was detected in our samples (Extended Data Table 2).

### Statistical analyses

Data were analysed using SAS® Studio University Edition version 3.8 to generate summary statistics and carry out analyses using generalized linear mixed model (GLMM) procedures. Models were generated by including all measured fixed effects and their interactions and the random effect of block. Where the statistical output from SAS indicated that one or more of the variance components were estimated to be zero, the random effect of block was removed, and the model was re-analyzed^59^. Although measures were repeated over two years in the same hoop houses, the population within the hoop houses changed from eight foundresses in 2017 to the 2nd generation of offspring including males and females in 2018. As such, year was considered a categorical effect rather than a repeated measure. The model with the lowest Akaike Information Criterion (AIC) was chosen as the best fit and statistical analyses were executed using that model. We performed *post hoc* pairwise comparisons for all the significant categorical fixed effects using differences of least square means, and a Tukey-Kramer-adjusted p >|t| for multiple comparisons. The significance level used in all tests was α=0.05.

Data is available from authors upon reasonable request.

## Acknowledgements

We would like to thank Katie Fisher, Dillon Muldoon, Loran Moran, Emma Jane Woods, Leonard Zettler, Matt Gibson, and especially Beatrice Chan for their excellent assistance in the field. This work was supported by Ontario Ministry of Agriculture, Food and Rural Affairs (OMAFRA) grant UofG2015-2466 (awarded to N.E.R. and D.S.W.C.), the Ontario Ministry of Environment and Climate Change (MOECC) Best in Science grant BIS201617-06 (awarded to N.E.R.), Natural Sciences and Engineering Research Council (NSERC) Discovery grants 2015-06783 (awarded to N.E.R.), the Fresh Vegetable Growers of Ontario (FVGO: awarded to N.E.R. and D.S.W.C.). D.S.W.C. was supported by the George and Lois Whetham Scholarship in Food Systems, an Ontario Graduate Fellowship, the Keith and June Laver Scholarship in Horticulture, the Fred W. Presant Scholarship, and a Latournelle Travel Scholarship. N.E.R. is supported as the Rebanks Family Chair in Pollinator Conservation by The W. Garfield Weston Foundation.

## Author Contributions

D.S.W.C. carried out the experimental work and subsequent data analyses. D.S.W.C. and N.E.R. conceived and designed the project and wrote the paper.

## Additional Information

**Supplementary information** is available

