## Extended data for "Neonicotinoid exposure affects foraging, nesting, and reproductive success of ground-nesting solitary bees"

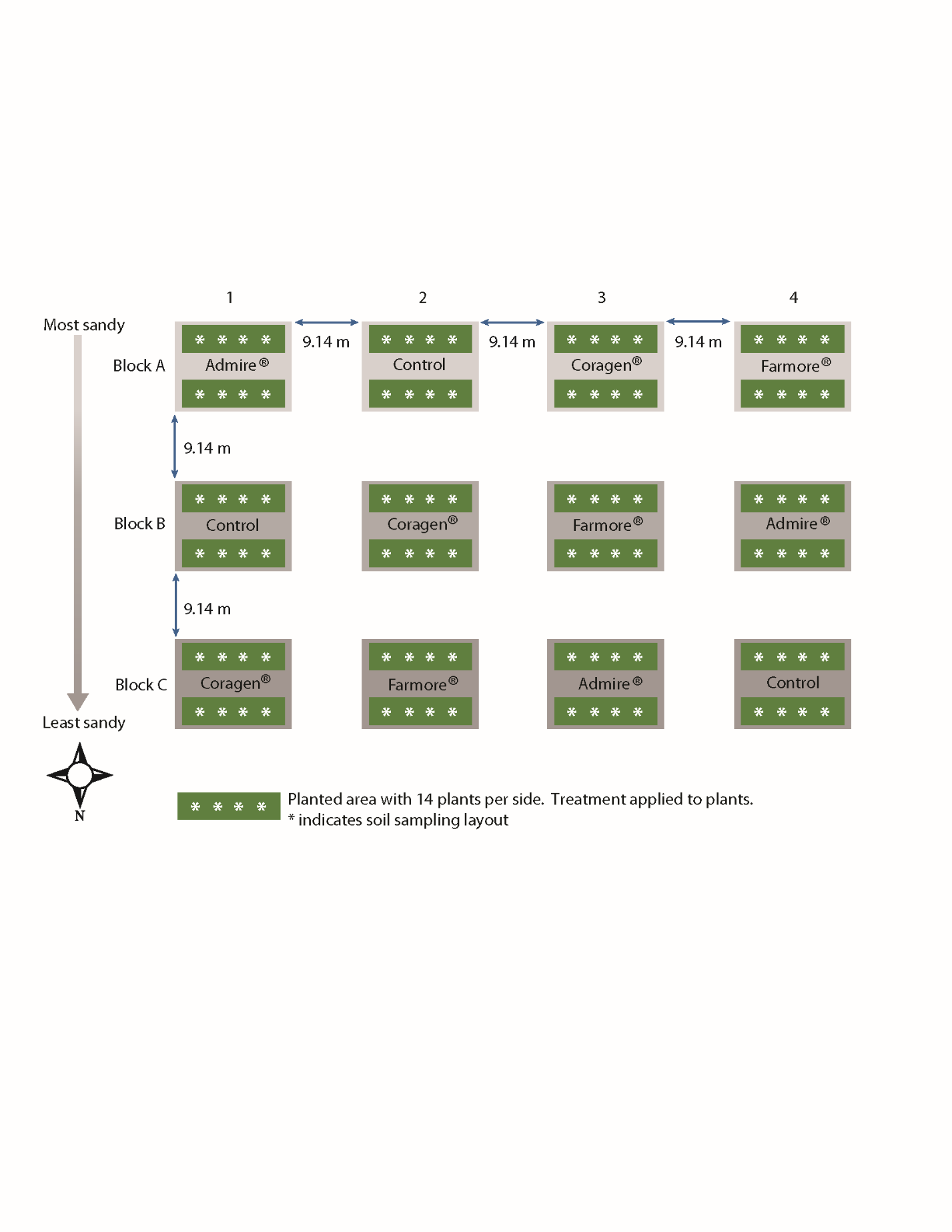

Extended Data Figure 1 Layout of study site in Peterborough County, Ontario, showing the arrangement and spacing of 12 hoop houses (grey rectangles: A1-C4), the treatments applied to each one (Admire®, Coragen® or FarMoreFI400®--described as FarMore®, or untreated control), the two planted areas in each hoop house (green rectangles), the soil sampling layout (white asterisks), and the soil texture gradient from most to least sandy at the site.

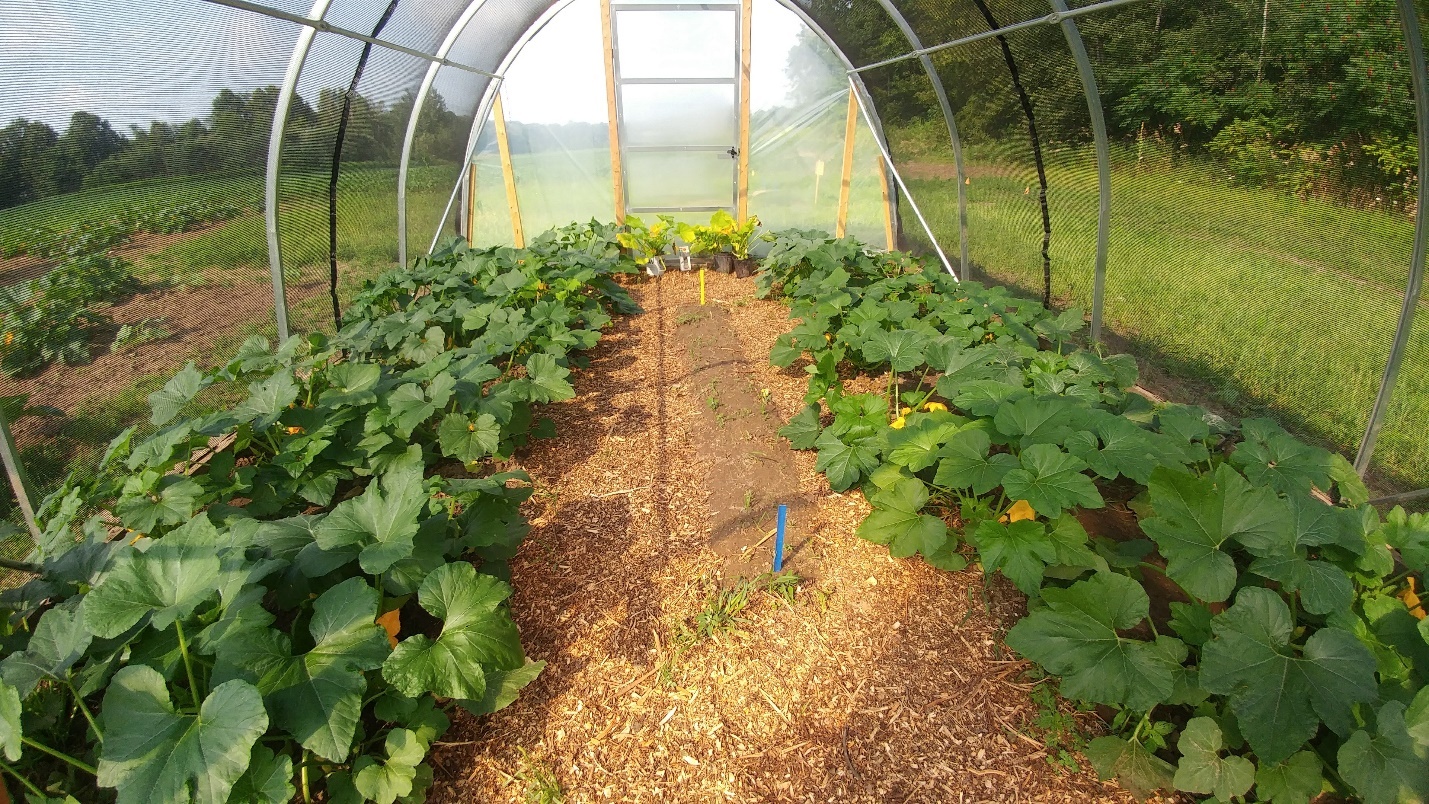

Extended Data Figure 2 Hoop house at the study site in Peterborough County, Ontario, showing the areas planted with 28 acorn squash plants, the mulched pathways, and the bare nesting area (between the blue and yellow vertical markers). Hoop houses were covered with shade cloth allowing exterior conditions to prevail inside while preventing introduced hoary squash bees from escaping or other bees from entering. Hoary squash bees used both the mulched paths and the bare soil area to excavate nests. Photo Beatrice Chan, used with permission.

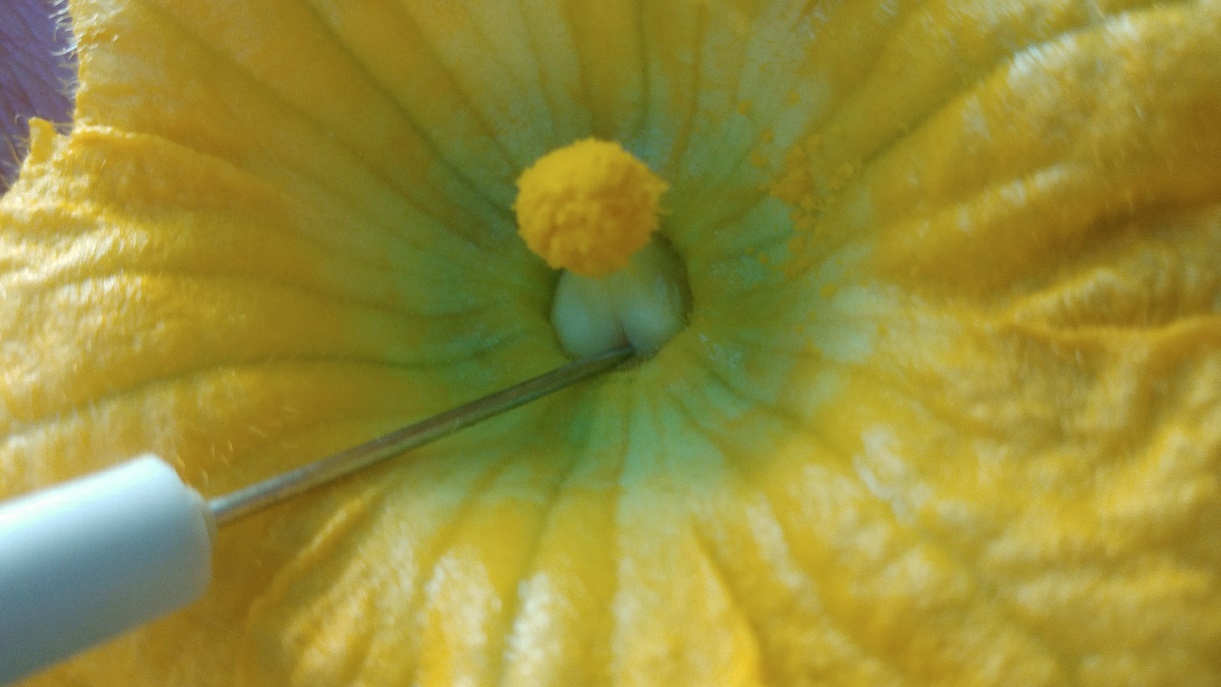

Extended Data Figure 3 Staminate flower of Table Star® variety of acorn squash. These flowers lacked access holes in the base of their fused stamen to allow bees to gather nectar from the enclosed nectaries. To solve this problem observers inserted a dissection needle into the base of the fused stamen to create three access holes for each staminate flower at the start of each day during the flowering period in 2017. Photo Beatrice Chan, used with permission.

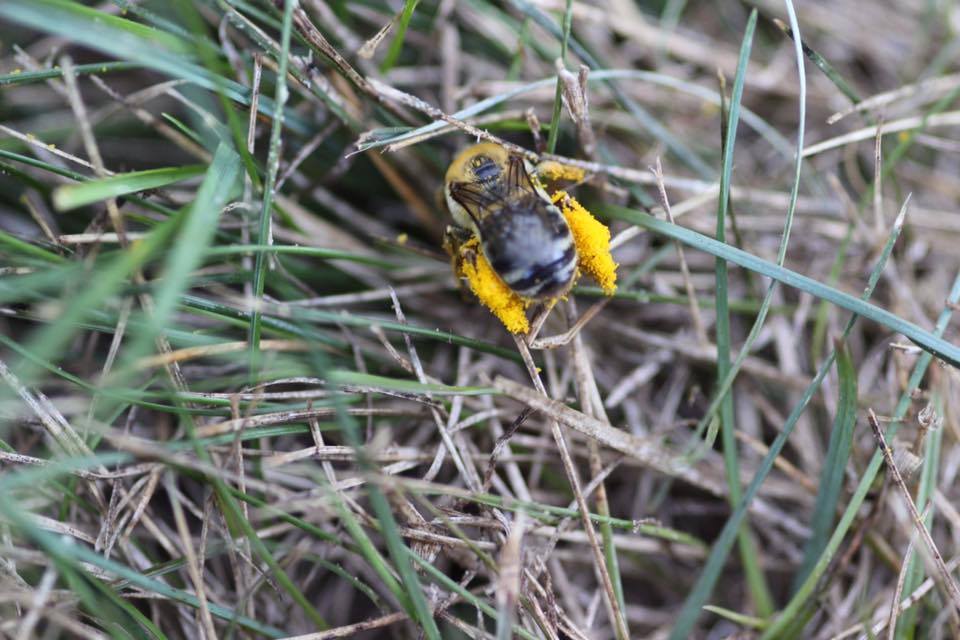

Extended Data Figure 4 Hoary squash bee female (*Eucera* (*Peponapis*) *pruinosa*) preparing to enter her nest with a full load of yellow *Cucurbita* spp. pollen on her hind legs. Female bees, such as the one pictured here, were captured for the study as they entered their nests. Photo Beatrice Chan, used with permission.

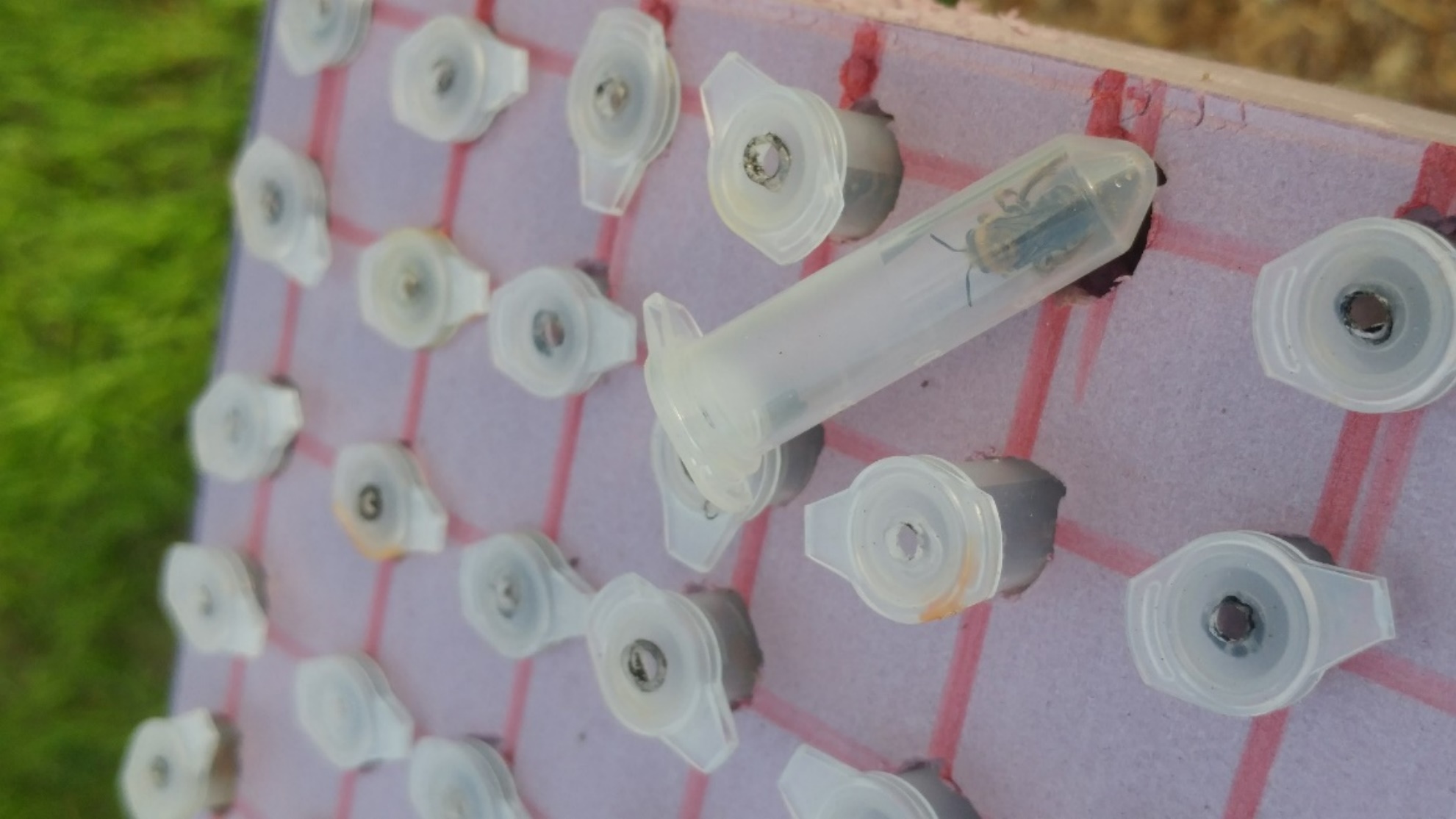

**Extended Data Figure 5 Aerated micro-centrifuge tubes used to contain and transport hoary squash bees (*Eucera* (*Peponapis*) *pruinosa*) from the site of capture in Guelph, Ontario to the study hoop houses in Peterborough County. Photo Beatrice Chan, used with permission.**

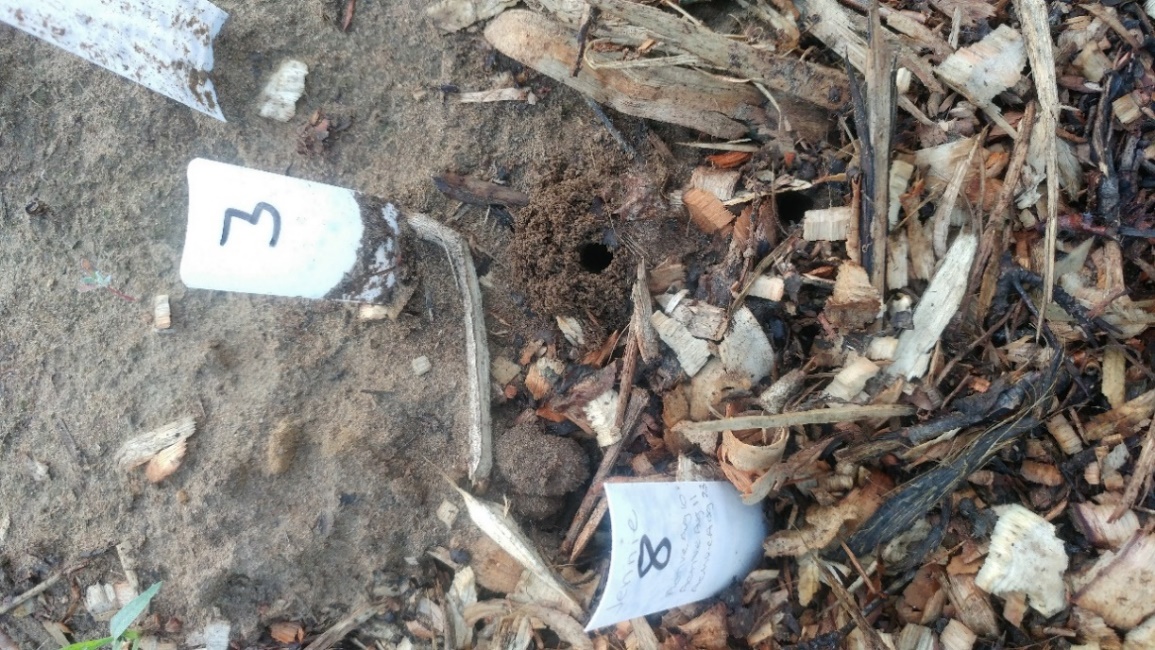

**Extended Data Figure 6 Aggregation of hoary squash bee nests within a study hoop house showing a nest entrance with tumulus of excavated soil (marked with a white arrow), and markers with identification numbers. These markers were used to enable observers to keep track of nests over the season and accurately record activity. Photo Beatrice Chan, used with permission.**

Extended Data Table 1 Sampling regime for each study including the year of observation, the number of observation days and the observation dates for the nest establishment, flower counts, and pollen harvest studies undertaken.

| **Study** | **Description** | **Year** | **Observation** | |
| --- | --- | --- | --- | --- |
|  |  |  | **Days** | **Dates** |
| Nest Establishment | Observers searched for active nests within each hoop house at the start of each observation day and marked each nest with a numbered and dated marker (Extended Data Fig.6) | 2017 | 10 | August 4,7,10,11,15,17,18,21,23,24 |
|  |  | 2018 | 8 | August 7,9,10,13,14,15,20,23 |
| Flower Counts | The total number of staminate and pistillate flowers in each hoop house was counted during the daily flowering period | 2017 | 6 | August 7,11,15,18,21,23 |
|  |  | 2018 | 8 | August 7,9,10,13,14,15,20,23 |
| Pollen Harvest | Pollen harvest by the whole population of hoary squash bees within a hoop house was evaluated by measuring the amount of unharvested pollen remaining on anthers of staminate flowers in each hoop house at the end of the daily foraging period | 2017 | 2 | August 17, 23 |
| Fruit set | Female flowers were marked each day for 10 days during the squash crop flowering period. At the end of the season, the number of flowers that had set fruit were counted and related to the total number of flowers | 2017 | 10 | August 4,7,10,11,15,17,18,21,23,24 |

**Extended Data Table 2 Mean concentration (ppb) of residues in soil from hoop houses treated with Admire® (applied to soil at planting) or FarMoreFI400® (applied as a seed coating) or Coragen® (spray applied to foliage at the 5-leaf stage) or an untreated control. All treatments were applied to squash (2017: Table Star®; 2018: Celebration®) in net-covered hoop houses before hoary squash bees were active. The mean concentration of residues in cells marked with a dash were below the minimum quantifiable or detectable limits. Six samples were taken in each treatment (i.e. 2 samples on each of 3 days) during the bee-active period in 2017 (July 17, August 4, August 18) and 2018 (July 18, August 15, August 23).**

|  | Hoop | Mean Residue Concentration Detected (ppb) | | | | | | | |
| --- | --- | --- | --- | --- | --- | --- | --- | --- | --- |
|  | House | imidacloprid | | clothianidin | | thiamethoxam | | chlorantraniliprole | |
| Treatment | ID | 2017 | 2018 | 2017 | 2018 | 2017 | 2018 | 2017 | 2018 |
| Admire® | A1 | 25.6 | 142.6 | 0.1 | - | - | - | - | - |
| Control | A2 | - | - | 0.2 | - | - | - | - | - |
| Coragen® | A3 | - | - | 1.0 | 1.3 | - | - | 0.6 | 11.3 |
| FarMoreF1400® | A4 | - | - | 1.2 | 1.2 | 0.1 | 1.9 | - | - |
| Control | B1 | - | - | 3.6 | - | 9.1 | - | - | - |
| Coragen® | B2 | - | - | 2.4 | - | - | - | - | 8.1 |
| FarMoreF1400® | B3 | - | - | 3.1 | 3.6 | - | 16.6 | - | - |
| Admire® | B4 | 11.3 | 44.6 | 1.7 | - | - | - | - | - |
| Coragen® | C1 | - | - | 5.7 | - | - | - | - | 23.8 |
| FarMoreF1400® | C2 | 31.5 | - | 2.6 | 1.6 | - | 1.1 | - | - |
| Admire® | C3 | 48.0 | 88.8 | 3.5 | 1.0 | - | - | - | - |
| Control | C4 | - | - | 3.6 | 2.2 | - | - | - | - |

Extended Data Figure 1 Layout of study site in Peterborough County, Ontario, showing the arrangement and spacing of 12 hoop houses (grey rectangles: A1-C4), the treatments applied to each one (Admire®, Coragen® or FarMoreFI400®--described as FarMore®, or untreated control), the two planted areas in each hoop house (green rectangles), the soil sampling layout (white asterisks), and the soil texture gradient from most to least sandy at the site.

Extended Data Figure 2 Hoop house at the study site in Peterborough County, Ontario, showing the areas planted with 28 acorn squash plants, the mulched pathways, and the bare nesting area (between the blue and yellow vertical markers). Hoop houses were covered with shade cloth allowing exterior conditions to prevail inside while preventing introduced hoary squash bees from escaping or other bees from entering. Hoary squash bees used both the mulched paths and the bare soil area to excavate nests. Photo Beatrice Chan, used with permission.

Extended Data Figure 3 Staminate flower of Table Star® variety of acorn squash. These flowers lacked access holes in the base of their fused stamen to allow bees to gather nectar from the enclosed nectaries. To solve this problem observers inserted a dissection needle into the base of the fused stamen to create three access holes for each staminate flower at the start of each day during the flowering period in 2017. Photo Beatrice Chan, used with permission.

Extended Data Figure 4 Hoary squash bee female (*Eucera* (*Peponapis*) *pruinosa*) preparing to enter her nest with a full load of yellow *Cucurbita* spp. pollen on her hind legs. Female bees, such as the one pictured here, were captured for the study as they entered their nests. Photo Beatrice Chan, used with permission.

**Extended Data Figure 5** Aerated micro-centrifuge tubes used to contain and transport hoary squash bees (*Eucera* (*Peponapis*) *pruinosa*) from the site of capture in Guelph, Ontario to the study hoop houses in Peterborough County. Photo Beatrice Chan, used with permission.

**Extended Data Figure 6** Aggregation of hoary squash bee nests within a study hoop house showing a nest entrance with tumulus of excavated soil (marked with a white arrow), and markers with identification numbers. These markers were used to enable observers to keep track of nests over the season and accurately record activity. Photo Beatrice Chan, used with permission.
