## Supplementary Information for "Neonicotinoid exposure affects foraging, nesting, and reproductive success of ground-nesting solitary bees"

Table S1 Model comparisons including descriptions and associated Akaike Information Criteria (AIC) for each model. Treatment effects refer to systemic insecticide treatment applied (Admire®, FarMore F1400® or Coragen®) or an untreated control. Flower effects refer to the number of flowers per hoop house in which the treatment was applied. Where data were collected in both 2017 and 2018, then year was included as a fixed effect in the model. Models for each dependent variable with lowest AIC values (shown in bold) were used for statistical analyses.

| **Dependent Variable** | **Model Iteration** | **Model Description** | **AIC** |
| --- | --- | --- | --- |
| Total Nests Initiated | 1 | Total Nests = Treatment + Year + Treatment*Year + Flowers, Random effect of Block | 104.3 |
|  | **2** | **Total Nests = Treatment + Year + Treatment*Year, Random effect of Block** | **101.8** |
|  | 3 | Total Nests = Treatment + Year + Flowers, Random effect of Block | 119.5 |
|  | 4 | Total Nests = Treatment + Year, Random effect of Block | 117.4 |
| Unharvested Pollen | **1** | **Unharvested Pollen = Treatment, Repeated effect of Observation Day, Random effect of Block** | **2307** |
| Total Offspring | **1** | **Total Offspring = Treatment + Year + Treatment*Year + Flowers, Random effect of Block** | **132.5** |
|  | 2 | Total Offspring = Treatment + Year + Treatment*Year, Random effect of Block | 163.5 |
|  | 3 | Total Offspring = Treatment + Year + Flowers, Random effect of Block | 163.5 |
|  | 4 | Total Offspring = Treatment + Year, Random effect of Block | 167.7 |
| Sex Ratio | 1 | Sex Ratio = Treatment + Year + Treatment*Year +Flowers, Random effect of block | 80.9 |
|  | **2** | **Sex Ratio = Treatment + Year** | 76.7 |
| % Fruit Set | **1** | **% Fruit Set = Treatment** | **3.2** |
| % Marketable Fruit | **1** | **% Marketable Fruit=Treatment + Year + Treatment*Year, Random effect of Block** | **-14.1** |

Table S2 Test statistics for each fixed effect associated with the best fitting model (Table S1) for each dependent variable (total nests initiated, total offspring, sex ratio, unharvested pollen, percentage fruit set, and percentage marketable fruit). If an effect was not included in the best fitting model this is indicated by “n/a”.

| **Model** | **Fixed Effects** | | | |
| --- | --- | --- | --- | --- |
|  | **Treatment** | **Year** | **Treatment*Year** | **Number of Flowers** |
| **Total Nests Initiated**  = Treatment + Year + Treatment*Year, Random effect of Block | F_(3,14)_ = 7.33  p = 0.0034 | F_(1,14)_ = 6.75  p = 0.0210 | F_(3,14)_ = 0.67  p = 0.5833 | n/a |
| **Unharvested Pollen** = Treatment, Repeated effect of Observation Day, Random effect of Block | F_(3,114)_ = 37.82  p < 0.0001 | n/a | n/a | n/a |
| **Total Offspring** = Treatment + Year + Treatment*Year + Flowers, Random effect of Block | F_(3,13)_ = 14.19  p = 0.0002 | F_(1,13)_ = 11.79  p = 0.0044 | F_(3,13)_ = 3.81  p = 0.0368 | F_(1,13)_ = 10.44  p = 0.0066 |
| **Sex Ratio** = Treatment + Year | F(_3,14)_ = 0.08  p = 0.9679 | F_(1,14)_ = 6.73  p = 0.0212 | n/a | n/a |
| **% Fruit Set** = Treatment | F_(3,8)_ = 0.16  p = 0.9194 | n/a | n/a | n/a |
| **% Marketable Fruit** = Treatment + Year + Treatment*Year, Random effect of Block | F_(3,14)_ = 0.14  p = 0.9360 | F_(1,14)_ = 9.22  p = 0.0089 | F_(3,14)_ = 1.17  p = 0.3547 | n/a |

Table S3 Post-hoc pairwise comparisons between treatments (Admire®, or Coragen®, or FarMore F1400®) and untreated control for dependent variables (total nests, total offspring, unharvested pollen) showing significant effects in best fit models (see Table S2). Tukey-Kramer adjustments were applied to p-values to account for multiple pairwise comparisons. Identical t-values for total nest comparisons reflect identical mean values of control, Coragen®, and FarMore F1400® treatments over both years (see Table 1). Lower and upper 95% confidence intervals are provided.

| **Dependent Variable** | **Treatments Compared** | **DF** | **t-Value** | **p > \|t\|** | **Adj p** |
| --- | --- | --- | --- | --- | --- |
| **Total Nests** | Admire® vs Control | 13 | -3.83 | 0.0018 | 0.0088 |
|  | Admire® vs Coragen® | 13 | -3.66 | 0.0029 | 0.0088 |
|  | Admire® vs FarMore F1400® | 13 | -3.68 | 0.0028 | 0.0088 |
|  | Control vs Coragen® | 13 | -0.00 | 0.9880 | > 0.9999 |
|  | Control vs FarMore F1400® | 13 | -0.00 | 0.9984 | > 0.9999 |
|  | Coragen® vs FarMore F1400® | 13 | -0.00 | 0.9996 | > 0.9999 |
| **Total Offspring** | Admire® vs Control | 15 | -5.98 | <0.0001 | 0.0002 |
|  | Admire® vs Coragen® | 15 | -5.33 | 0.0001 | 0.0007 |
|  | Admire® vs FarMore F1400® | 15 | -3.55 | 0.0035 | 0.0163 |
|  | Control vs Coragen® | 15 | 0.80 | 0.4393 | 0.8542 |
|  | Control vs FarMore F1400® | 15 | 2.59 | 0.0223 | 0.0910 |
|  | Coragen® vs FarMore F1400® | 15 | 1.82 | 0.0924 | 0.3094 |
| **Unharvested Pollen** | Admire® vs Control | 114 | 9.38 | <0.0001 | <0.0001 |
|  | Admire® vs Coragen® | 114 | 6.72 | <0.0001 | <0.0001 |
|  | Admire® vs FarMore F1400® | 114 | 9.02 | <0.0001 | <0.0001 |
|  | Control vs Coragen® | 114 | -2.65 | 0.0091 | 0.0443 |
|  | Control vs FarMore F1400® | 114 | -0.36 | 0.7215 | 0.9843 |
|  | Coragen® vs FarMore F1400® | 114 | 2.30 | 0.0234 | 0.1047 |

Table S4 Raw data for flower count, total reproductive females, total nests, total offspring, male offspring, and female offspring and calculated indices (percentage of male, and sex ratio of, offspring produced) for each pesticide treatment applied to a squash crop in twelve net-covered hoop houses occupied by a captive population of hoary squash bees (*Eucera* (*Peponapis*) *pruinosa*).

| **Treatment** | **Hoop House** | **Year** | **Mean Flower Count** | **Number of Reproductive Female Bees** | **Total Nests** | **Total Offspring** | **Male Offspring** | **Female Offspring** | **Percent Males (%)** | **Sex Ratio** |
| --- | --- | --- | --- | --- | --- | --- | --- | --- | --- | --- |
| Admire® | 1A | 2017 | 41.17 | 8 | 0 | 3 | 1 | 2 | 33 | 0.50 |
| Admire® | 3C | 2017 | 43.00 | 8 | 1 | 6 | 2 | 4 | 33 | 0.50 |
| Admire® | 4B | 2017 | 52.17 | 8 | 0 | 1 | 1 | 0 | 100 | - |
| Admire® | 1A | 2018 | 56.00 | 2 | 0 | 0 | 0 | 0 | - | n/a |
| Admire® | 3C | 2018 | 53.00 | 4 | 8 | 10 | 8 | 2 | 80 | 4.00 |
| Admire® | 4B | 2018 | 54.25 | 0 | 0 | 0 | 0 | 0 | - | n/a |
| Control | 1B | 2017 | 44.50 | 8 | 13 | 22 | 13 | 9 | 59 | 1.44 |
| Control | 2A | 2017 | 27.83 | 8 | 4 | 5 | 3 | 2 | 60 | 1.50 |
| Control | 4C | 2017 | 56.17 | 8 | 9 | 42 | 15 | 27 | 36 | 0.56 |
| Control | 1B | 2018 | 54.13 | 9 | 8 | 58 | 44 | 14 | 76 | 3.14 |
| Control | 2A | 2018 | 43.63 | 2 | 6 | 31 | 19 | 12 | 61 | 1.58 |
| Control | 4C | 2018 | 41.63 | 27 | 19 | 37 | 25 | 12 | 68 | 2.08 |
| Coragen® | 1C | 2017 | 57.33 | 8 | 9 | 17 | 5 | 12 | 29 | 0.42 |
| Coragen® | 2B | 2017 | 41.00 | 8 | 8 | 13 | 6 | 7 | 46 | 0.86 |
| Coragen® | 3A | 2017 | 45.83 | 8 | 7 | 24 | 5 | 19 | 21 | 0.26 |
| Coragen® | 1C | 2018 | 57.63 | 12 | 11 | 62 | 35 | 27 | 56 | 1.30 |
| Coragen® | 2B | 2018 | 46.25 | 7 | 10 | 49 | 45 | 4 | 92 | 11.25 |
| Coragen® | 3A | 2018 | 32.38 | 12 | 14 | 17 | 7 | 10 | 41 | 0.70 |
| FarMore FI400® | 2C | 2017 | 63.67 | 8 | 8 | 30 | 9 | 21 | 30 | 0.43 |
| FarMore FI400® | 3B | 2017 | 49.00 | 8 | 8 | 17 | 10 | 7 | 59 | 1.43 |
| FarMore FI400® | 4A | 2017 | 51.17 | 8 | 2 | 18 | 8 | 10 | 44 | 0.80 |
| FarMore FI400® | 2C | 2018 | 41.13 | 21 | 13 | 34 | 30 | 4 | 88 | 7.50 |
| FarMore FI400® | 3B | 2018 | 34.63 | 6 | 10 | 15 | 9 | 6 | 60 | 1.50 |
| FarMore FI400® | 4A | 2018 | 44.88 | 10 | 18 | 11 | 8 | 3 | 73 | 2.67 |
